# Nash equilibria in human sensorimotor interactions explained by Q-Learning

**DOI:** 10.1101/2021.06.14.448333

**Authors:** Cecilia Lindig-León, Gerrit Schmid, Daniel A. Braun

**Author notes:** these authors contributed equally to this work (Cecilia Lindig-León, Gerrit Schmid).

## Abstract

The Nash equilibrium concept has previously been shown to be an important tool to understand human sensorimotor interactions, where different actors vie for minimizing their respective effort while engaging in a multi-agent motor task. However, it is not clear how such equilibria are reached. Here, we compare different reinforcement learning models based on haptic feedback to human behavior in sensorimotor versions of three classic games, including the Prisoners’ Dilemma, and the symmetric and asymmetric matching pennies games. We find that a discrete analysis that reduces the continuous sensorimotor interaction to binary choices as in classical matrix games does not allow to distinguish between the different learning algorithms, but that a more detailed continuous analysis with continuous formulations of the learning algorithms and the game-theoretic solutions affords different predictions. In particular, we find that Q-learning with intrinsic costs that disfavor deviations from average behavior explains the observed data best, even though all learning algorithms equally converge to admissible Nash equilibrium solutions. We therefore conclude that it is important to study different learning algorithms for understanding sensorimotor interactions, as such behavior cannot be inferred from a game-theoretic analysis alone, that simply focuses on the Nash equilibrium concept, as different learning algorithms impose preferences on the set of possible equilibrium solutions due to the inherent learning dynamics.

## Introduction

Nash equilibria are the central solution concept for understanding strategic interactions between different agents^1^. Crucially, unlike other maximum expected utility decision-making models^2,3^, the Nash equilibrium concept cannot assume a static environment that can be exploited to find the optimal action in a single sweep, but it rather defines a fixed point representing a combination of strategies that can be found by iteration, so that finally no agent has anything to gain by deviating from their equilibrium behaviour^4,5^. Here, a strategy is conceived as a probability distribution over actions, so that Nash equilibria are in general determined by combinations of probability distributions over actions (mixed Nash equilibria), and only in special cases by combinations of single actions (pure equilibria)^6,7^. An example of the first kind is the popular rock-papers-scissors game which can be simplified to the matching pennies game with two action choices, where in either case the mixed Nash equilibrium requires players to randomize their choices uniformly. An example of the second kind is the prisoners’ dilemma where both players choose between cooperating and defecting without communication. The pure Nash equilibrium in this game requires both players to defect, because the payoffs are designed in a way that allow for a dominant strategy from the perspective of a single player, where it is always better to defect, no matter what the other player is doing.

The Nash equilibrium concept has not only been broadly applied in economic modeling of interacting rational agents, companies and markets^8,9^, but also to explain the dynamics of animal conflict^10,11^, population dynamics including microbial growth^12,13^, foraging behavior^14–17^, the emergence of theory of mind^18–21^, and even monkeys playing rock-papers-scissors^22,23^. Recently, Nash equilibria have also been proposed as a concept to understand human sensorimotor interactions^24–27^. In these studies human dyads are typically coupled haptically and experience physical forces that can be regarded as payoffs in a sensorimotor game. By designing the force payoffs appropriately in dependence of subjects’ actions, these sensorimotor games can be made to correspond to classic pen-and-paper games like the prisoner’s dilemma with a single pure equilibrium, coordination games with multiple equilibria, or signaling games with Bayesian Nash equilibria to model optimal sensorimotor communication. In general, it was found in these studies^28,29^ that subjects’ sensorimotor behavior during haptic interactions was in agreement with the game theoretic predictions of the Nash equilibrium, even though the pen-and-paper versions of some games systematically violate these predictions. This raises the question of how such equilibria are attained, especially since the Nash equilibrium concept itself provides no explanation of how it is reached, especially when there are multiple equivalent equilibria.

In this study we investigate what kind of learning models could explain how humans that are interacting through a haptic sensorimotor coupling reach a Nash equilibrium. The problem of learning in games can be approached within different frameworks, including learning of simple fixed response models like partial best response dynamics for reaching pure Nash equilibria^30,31^, or fictitious play with smoothed best response dynamics for mixed equilibria^32,33^, as well as more sophisticated reinforcement learning models like Q-learning^34,35^, policy gradients^36,37^, minimax Q-learning^38^ or Nash Q-learning^39,40^, together with learning models in evolutionary game theory for reaching Nash equilibria (evolutionary stable strategies) through population dynamics^41^. Here we focus on model-free reinforcement learning models to explain subjects’ sensorimotor interactions, as in many of the previous experiments^24,25,28^ subjects interact only through haptic feedback and cannot otherwise “see” the choices of the other player. In particular, we consider model-free reinforcement learning models like Q-learning and direct policy search methods like policy gradients that exclusively rely on force feedback during learning. We compare Q-learning and policy gradient learning in games with pure and mixed Nash equilibria, including the prisonners’ dilemma and two versions of the matching pennies game with symmetric and asymmetric pay-offs respectively. While the prisonners’ dilemma game has a single pure equilibrium, the sensorimotor version of the matching pennies games has infinitely many mixed Nash equilibria that are theoretically equivalent, and we investigate whether the different learning algorithms introduce additional preferences between these equilibrium strategies based on the inherent learning dynamics and we check how these match up with human learning behavior.

## Results

To investigate learning in pure and mixed equilibria during sensorimotor interactions, we designed three continuous sensorimotor games that are variants of the traditional two-player matrix games of the prisoner’s dilemma, the asymmetric matching pennies game, and the symmetric matching pennies game (see section 1). During the experiment two players were sitting next to each other and interacted through a virtual reality system with the handles of individual robotic interfaces that were free to move in the horizontal plane (see Figure 6A). In each trial, players were requested to move their handle to cross a target bar ahead of them, where the lateral position of both handles determined the individual magnitude of a resistive force opposing the forward motion of the handle. Thus, the movement of each player directly impacted on the forces experienced by both players in a continuous fashion. This way, our sensorimotor game differs in three essential aspects from the traditional cognitive versions, in that first, we have an implicit effort cost through haptic coupling, second, subjects could choose from a continuum of actions defining a continuous payoff landscape, where the payoffs in the four corners correspond to the payoffs in the traditional payoff matrices, and third, that subjects are unaware of the structure of the haptic coupling, i.e., they do not have complete information about the payoffs, as they only have access to their own payoffs through force feedback.

Since the payoffs can only be learned from experience, we devised different reinforcement learning models that can emulate subjects’ behavior by adapting their choice distributions in a way that avoids punitive forces. As subjects were only communicated their own payoff when making their choices, we compare only model-free reinforcement learning schemes, in particular Q-learning and policy gradient models where we consider two reward conditions, with and without intrinstic costs added to the extrinsically imposed reward defined by the haptic forces. In essence, the intrinsic costs make it more expensive for the learner to change actions dramatically between trials—see Methods for details. We focus on trial-by-trial learning across the 40 trial blocks where games were repeated and neglect within-trial adaptation, as we found that initial and final positions of movement trajectories were close together most of the time. In particular, we found that in approximately 65% of the trials for the prisoner’s dilemma, and in 90% and 95% of the trials for the asymmetric and symmetric matching pennies respectively, the subjects’ final decision laid within a 3.6cm neighborhood (20% of the entire workspace) of their initial position in each trial, and there was no systematic change across the block of trials. We found that the distribution of distances between the initial and final position in each trial did not change significantly when comparing the first 20 trials of each block with the last 20 trials of each block (Kolmogorov-Smirnov test, *p* > 0.05 for all games). Therefore we concentrate only on the final position of each trial, where the force pay-off was strongest.

### Categorical Analysis

Previously, we have studied subjects’ choices in the sensorimotor game in a binary fashion^24,28^ to allow for a direct comparison with the corresponding pen-and-paper 2-by-2 matrix game. To this end, we divide each subject’s action space into two halves and categorize actions accordingly—compare Figure 1. The four quadrants of the combined action space then reflect all possible combinations, for example in the prisoners’ dilemma (*defect, defect*), (*defect, cooperate*), (*cooperate, defect*) and (*cooperate, cooperate*). When comparing the histogram over subjects’ choices over the first 10 trials compared to the last 10 trials of each block of 40 trials, it can be clearly seen that behavior adapts and becomes more consistent with the Nash equilibrium solution of the 2-by-2 matrix games given by the prisoners’ dilemma and the asymmetric matching pennies game. In particular, in the prisoners’ dilemma game, subjects increasingly concentrate on the defect/defect solution, even though the Nash solution is never perfectly reached, as other action combinations also occur late in learning. In the asymmetric matching pennies game, player 1 displays the expected asymmetric action distribution, whereas player 2 shows the expected 50:50 behavior. In both games, the of the joint action distribution decreases significantly over the block of 40 trials—see Supplementary Figure 1. In the case of the symmetric matching pennies game, the histograms do not reveal any learning or entropy decrease, as expected, because the random Nash strategy coincides with the random initial strategy.

**Figure 1.**
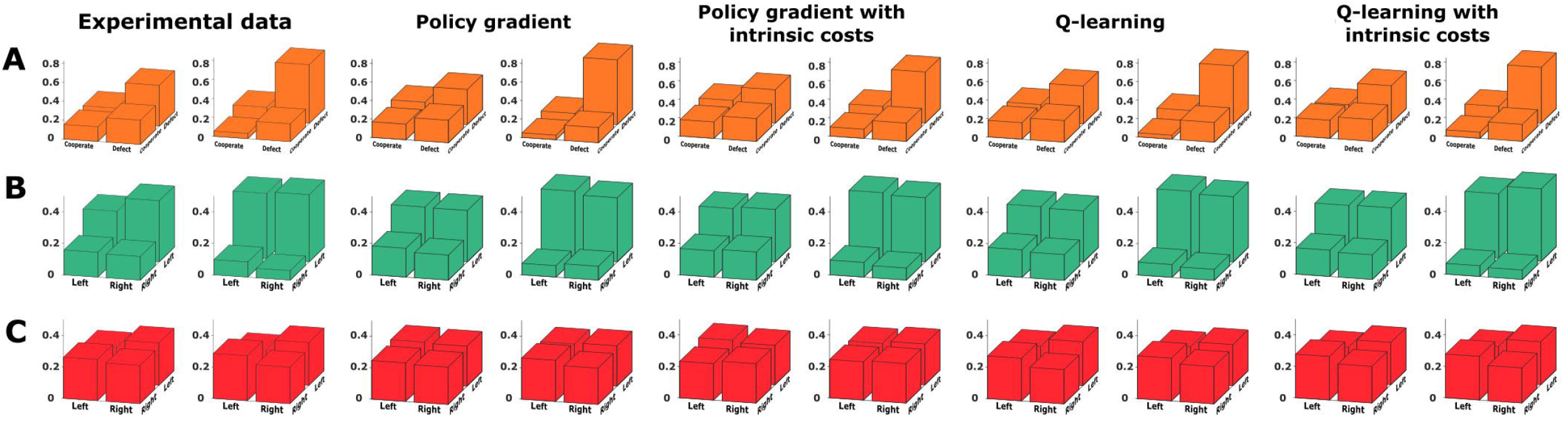
Categorical analysis. Final decisions in the *x*1*x*2-space categorized into two halves for the experimental data (left) and, from left to right, the results of the binary gradient descent, binary Q-learning, binary gradient descent with prior cost, and binary Q-learning with prior cost. We show the first 10 trials compared to the last 10 trials of each block of 40 trials.**(A) Prisoner’s dilemma. (B) Asymmetric matching pennies. (C) Matching pennies.**

We compared the binary *Q*-learning described in the methods as model 1, and the policy gradient described as model 2, to the human data by exposing both models to the same kind of game sequence that was experienced by the subjects. In the simulations, we always applied the same learning model for both players, albeit with player-specific parameters that were fitted to match subjects’ action distributions. In Figure 1 it can be seen that both kinds of learning models explain the subjects’ behavior equally well. Also when repeating the same analysis including the intrinstic cost function that discourages deviations from previous positions the result remains essentially the same. This leaves open the possibility that the categorical analysis may be too coarse to distinguish between these kinds of models. In the following, we therefore study the continuous games in terms of their continuous responses and not just their binary discretization. We compare subjects’ behavior in the three different games to two continuous reinforcement learning models, where one is a policy gradient model with a continuous action distribution, and the other one a continuous Q-learning model.

### Continuous Analysis

#### Prisoner’s dilemma

In Figure 2A subjects final decisions over the continuous Prisoners’ Dilemma game are shown as a scatter plot within the *x*_1_*x*_2_-space, where the single pure Nash equilibrium is located in the top-right corner at position (1, 1). Over the course of 40 trials subjects’ responses increasingly cluster around the Nash equilibrium, even though a considerable spread remains. The difference in endpoint distribution between the first 10 trials in a block and the last 10 trials was highly significant for player 1 and player 2 (KS-test, *p* < 0.01 for each player). A two-dimensional histogram binning of the experimental scatter plots can be found in Figure 2B.

**Figure 2.**
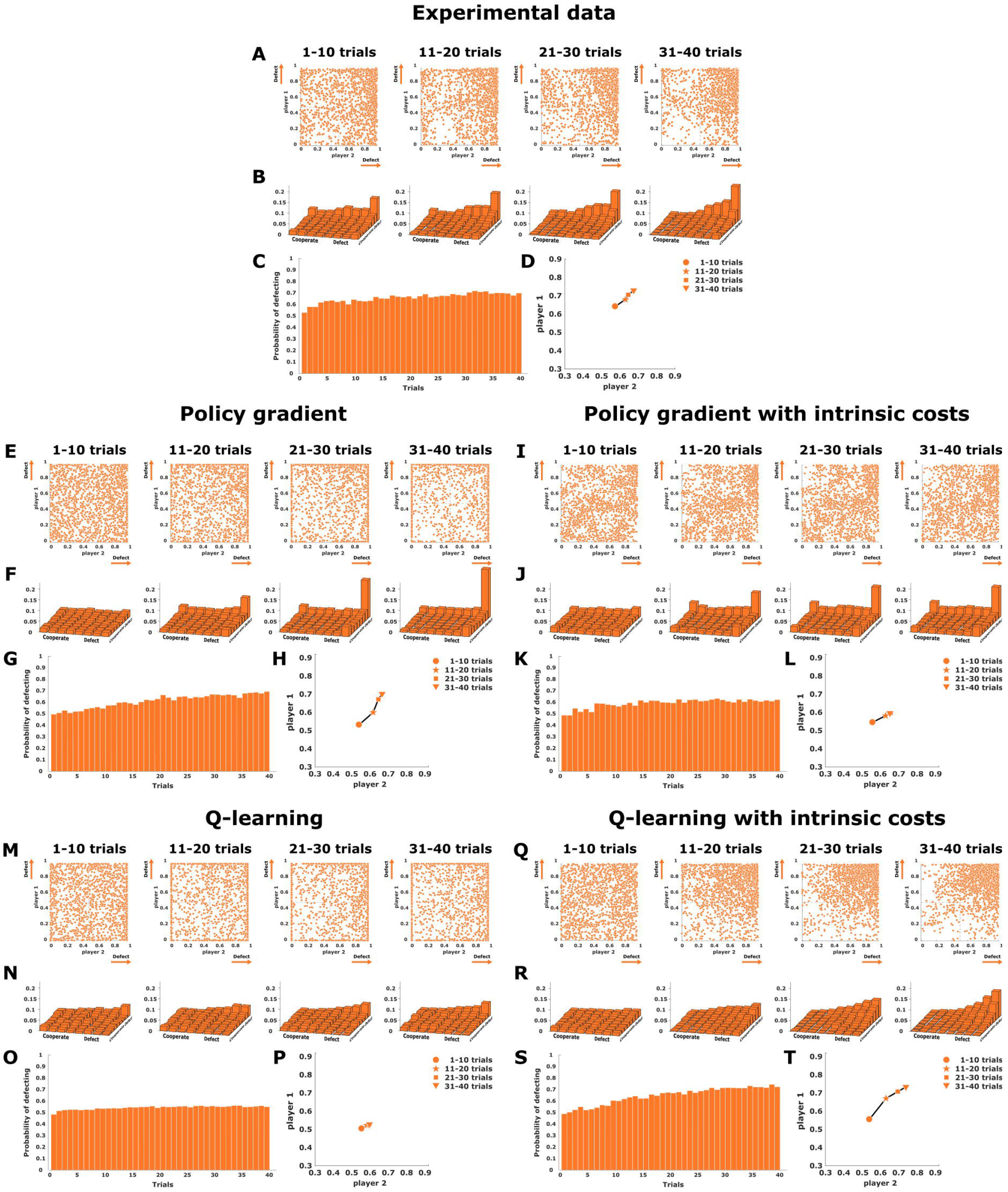
Prisoner’s dilemma. (A), (E), (I), (M), and (N) show scatter plots of final decisions in the *x*1*x*2-plane, where subjects’ actions are expected to cluster around the single pure Nash equilibrium located in the top-right corner at position (1,1). (B), (F), (J), (N), and (R) show two-dimensional histograms binning the experimental scatter plots. (C), (G), (K), (O), and (S) represent the change of the mean endpoints (averaged for both players) for each trial across the block of 40 trials. (D), (H), (L), (P), and (t) show the direction of adaptation in the endpoint space. The experimental data is shown at the top, the four continuous models are shown below.

Figure 2E shows the performance of two coupled continuous policy gradient learners on the same task. The model learners also converge to the Nash solution, but the variability is spread almost uniformly along the boundaries and corners, which is in stark contrast to subjects’ behavior. The same is also true if the policy gradient learners are parameterized slightly differently with the logit-normal—see Supplementary Figure 2. Figure 2M shows two coupled continuous Q-learning agents performing the prisoners’ dilemma sensorimotor task. The model learners slowly start concentrating probability mass in the quadrant of the Nash equilibrium, but their responses are more spread out than the subjects’ responses, giving in comparison a distribution that is too flat due to the broad exploration. One way to enforce more specific exploration is through the consideration of intrinsic costs, for example by assuming that selecting actions that significantly deviate from the action prior, given by averaging over all trials, is costly^42, 43^. Figure 2I and Figure 2Q show the performance of the two learning models considering such an intrinsic cost. While the behavior of the gradient learner still shows a sharp peak in the corner, the Q-learning model concentrates actions in a way that is more similar to human behavior in the quadrant of the Nash equilibrium, and it even shows the exponential decline in probability that is observed in the human data when moving away from the Nash corner. Accordingly, the Q-learning model with intrinsic costs achieves the most similar two-dimensional histogram to the data, as quantified by the Euclidean distance between the subjects’ and the models’ histograms—compare Supplementary Table 1.

A statistical measure to assess the effects of learning is the change of the mean endpoint (averaged for both players) for each trial across the block of 40 trials. Figure 2C shows a slow increase from 0.5 to approximately 0.7. The Q-learning model without intrinsic costs shows again a flat distribution resulting from the broad exploration (see Figure 2O). In contrast, both the gradient learning models (see Figures 2G and 2K) and the local Q-learning model with intrinsic costs (see Figure 2S), manage to mimic the experimental responses, with the latter model achieving the lowest mean-squared error compared to the data—compare Supplementary Table 1.

To study the direction of adaptation in the endpoint space, we look at the difference vectors resulting from subtracting the mean endpoints of each mini-block of 10 trials from the preceding mini-block of 10 trials. The arrow plot in Figure 2D shows that subjects move towards the Nash equilibrium on a straight path, where the step size of learning becomes smaller over time. Again this adaptation pattern is reproduced by both the gradient learning models ((see Figures 2H and 2L)) and by the Q-learning model with intrinsic costs (see Figure 2T). Without the intrinsic costs, the extensive search of the Q-learning agents generates a random adaptation (see Figure 2P).

In summary, our analysis demonstrates that it is not enough to look at the mean behavior to distinguish between the different models. In fact, when looking at the entire two-dimensional histogram over the continuous action space, it becomes apparent that the Q-learning model with intrinsic costs is the only one that can capture subjects’ behavior well, as it is the only model that concentrates actions in the Nash quadrant where the concentration increases gradually with proximity to the Nash solution (compare Figures 2B, 2F, 2J, 2N, and 2R).

#### Asymmetric matching pennies

Figure 3A shows subjects’ final decisions in the continuous asymmetric matching pennies game as a scatter plot in the *x*_1_*x*_2_-plane, where the set of mixed Nash equilibria corresponds to all joint distributions *p*(*x*_1_,*x*_2_) where the marginal distribution of player 1 has the expected location 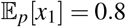 and the marginal distribution of player 2 has the expected location 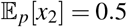. Throughout the course of 40 trials, subjects’ responses evolve from a symmetric distribution centered approximately at (0.5,0.5) to an asymmetric distribution that is tilted for player 1 as expected—compare Figure 3D. The difference in endpoint distribution between the first 10 trials in a block and the last 10 trials was highly significant (KS-test, *p* < 0.01). A two-dimensional histogram binning of the scatter plots can be seen in Figure 3B.

**Figure 3.**
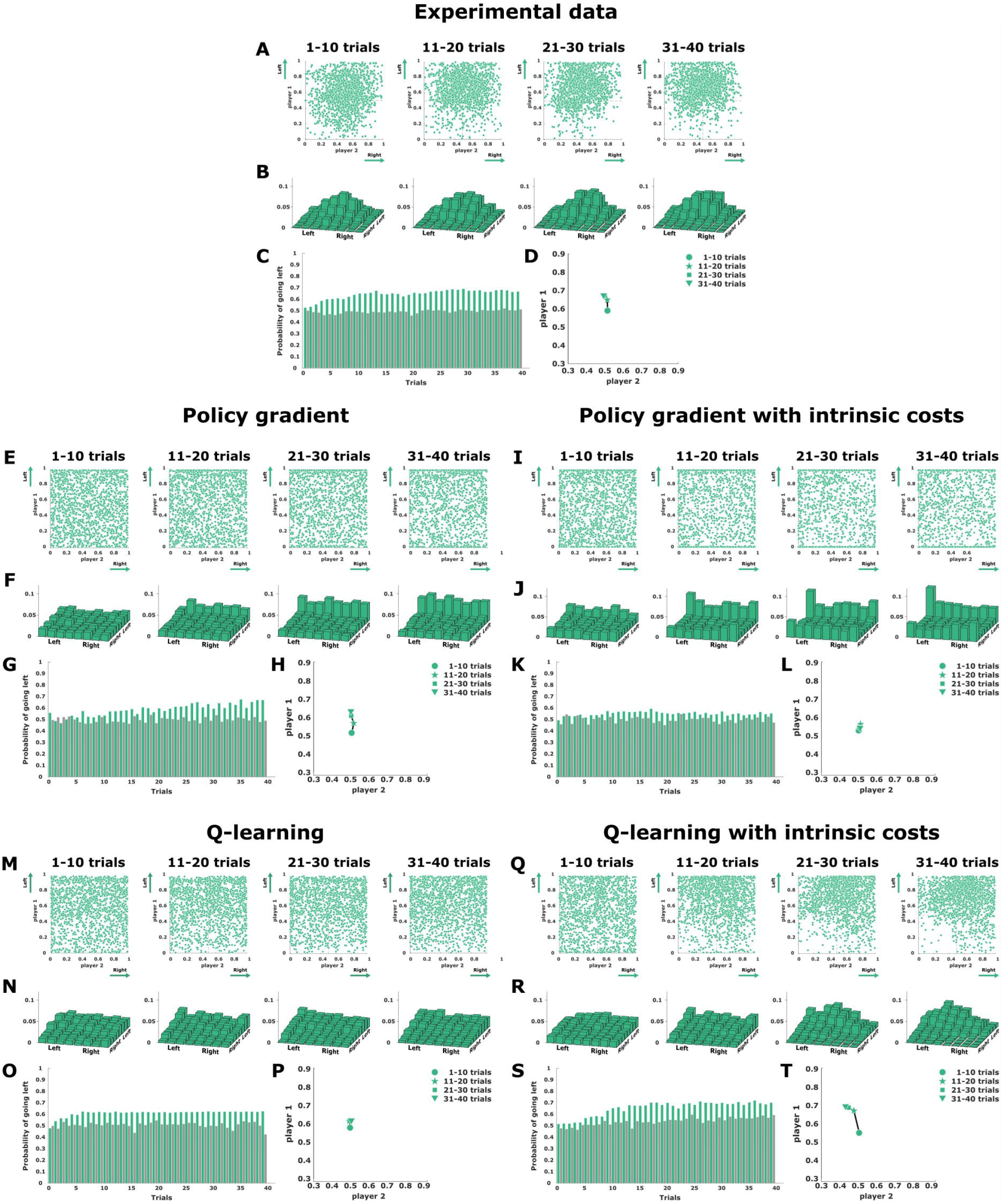
Asymmetric matching pennies. (A), (E), (I), (M), and (N) show final decisions as a scatter plot in the *x*1*x*2-plane, where subjects’ actions are expected to cluster in top quadrants along each mini-block of 10 trials. (B), (F), (J), (N), and (R) show a two-dimensional histogram binning of the experimental scatter plots. (C), (G), (K), (O), and (S) present the change over the mean endpoint (averaged for both players) for each trial across the block of 40 trials. (D), (H), (L), (P), and (t) show the direction of adaptation in the endpoint space. The experimental data is shown on the top, the four continuous models are below.

Figure 3E shows the endpoint distribution of two coupled policy gradient learners performing in the continuous asymmetric matching pennies game. As expected player 1 shifts to an asymmetric behavior, while player 2 remains random, but the model actions are mainly spread along the two opposing boundaries, which singles out a possible Nash solution, but is in stark contrast to subjects’ behavior that concentrates in the upper quadrants of the workspace. This preference for extreme responses of the gradient model is even more extreme when including intrinsic costs, which eliminates most of the asymmetry (see Figure 3I).

Figure 3M shows two coupled continuous Q-learning agents performing the same task. In contrast to the gradient learning models, the best-fitting Q-learning models manage to preserve substantial probability mass in the center of the workspace, but this happens at the expense of such a slow learning progress that the resulting action distribution is close to uniform, and therefore also does not capture subjects’ behavior well. This changes, however, when including intrinsic costs. Figure 3Q shows two fitted continuous Q-learning agents with intrinsic costs that manage to qualitatively capture the experimentally observed distribution in that player 1 concentrates its probability mass in the top quadrants, whereas player 2 shows more uniform behavior, as expected.

Figure 3C shows the change of the mean endpoint (independently for each player) in each trial across the block of 40 trials. We can observe an increase from 0.5 to approximately 0.8 for player 1, whereas for player 2 the mean value remains constant along all trials around 0.5. This behavior is also observed in Figure 3D, which shows the average direction of subjects’ adaptation. These adaptation patterns are properly reproduced by the gradient learning model (see Figures 3G and 3H), however, when including the intrinsic costs the symmetry between extreme responses causes a random behavior in both players (see Figures 3K and 3L). We see clearly that the Q-learning model without intrinsic costs depicted in Figure 3O is very slow in learning, which can also be seen from the mean direction of adaptation depicted in Figure 3P. However, when considering intrinsic costs in the Q-learning model, the change of the mean endpoint (see Figure 3S) and the average direction of adaptation (see Figure 3T) match subjects behavior quite well.

And even though the Q-learning model with intrinsic costs is initially slightly slower than the human subjects in picking up the equilibrium behavior, taken together with the two-dimensional histogram binning of the scatter plots in Figures 3B, 3F, 3J, 3N, 3R the Q-learning model with intrinsic costs provides the best fit to subjects’ behavior, confirmed by the lowest Euclidean distance between model and subject histograms—compare Supplementary Table 1.

#### Symmetric matching pennies

While the asymmetric matching pennies game requires adaptation in action frequency, learning in the symmetric game is not expected to show changes in the action frequency, and thus we finally need to look for other signatures of learning. In Figure 4A subjects final decisions in the continuous matching pennies game are shown as a scatter plot in the *x*_1_*x*_2_-plane, where the set of mixed Nash equilibria corresponds to all joint distributions *p*(*x*_1_, *x*_2_) where the marginal distribution of player 1 and player 2 has the expected location 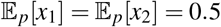. Subjects’ responses generally cluster around the center position with mean (0.5, 0.5) in the first block of 10 trials and remain that way through the entire block of 40 trials. A two-dimensional histogram binning of the scatter plots can be seen in Figure 4B. While subjects’ initial behavior can be explained as a result of ignorance, the same behavior is also compatible with the final Nash behavior, and so the response frequencies remain stable throughout the block of 40 trials. This can be seen from the mean endpoint (averaged for both players) across the block of 40 trials shown in Figure 4C, and from the average direction of subjects’ adaptation depicted in Figure 4D.

**Figure 4.**
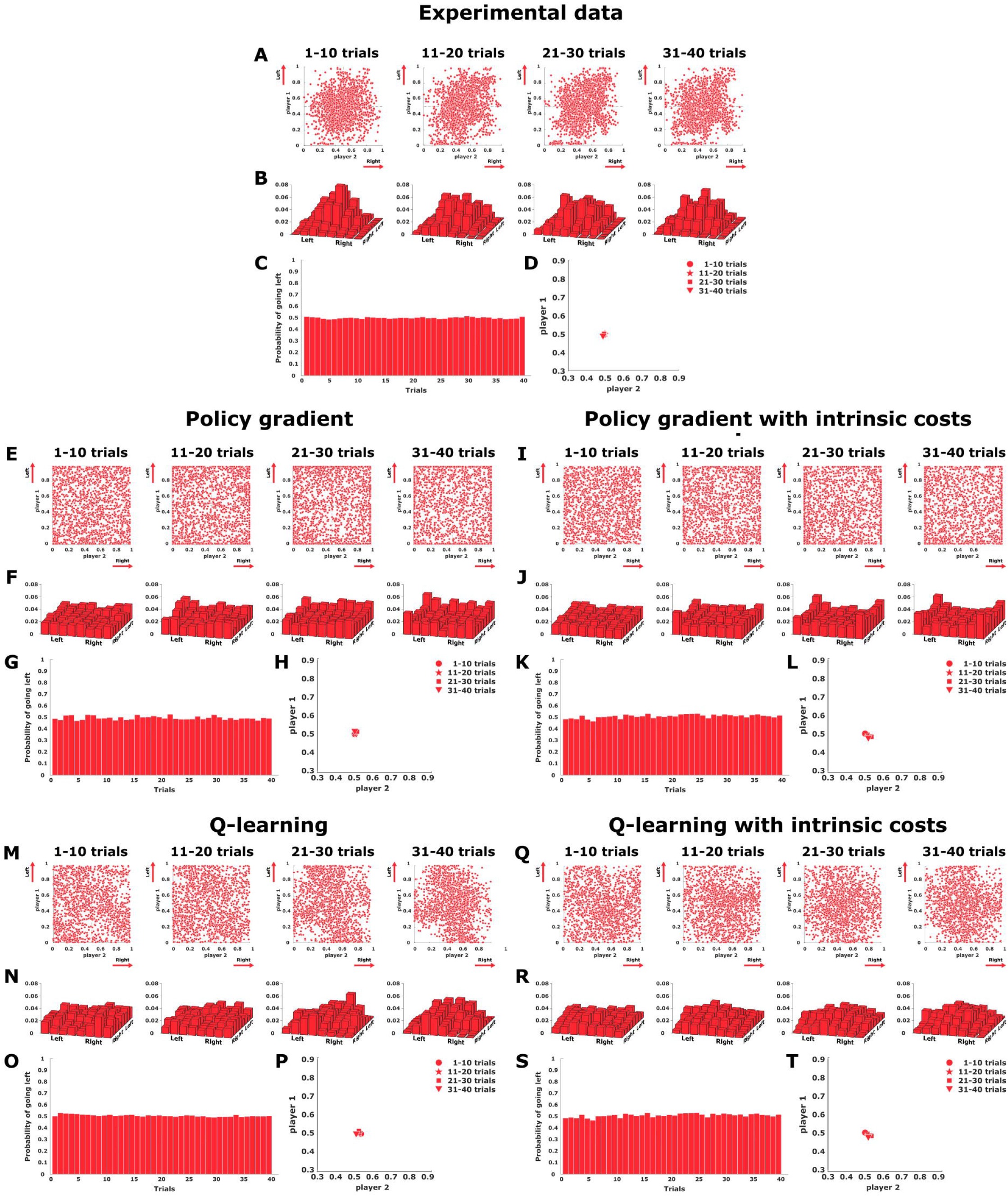
Symmetric matching pennies. (A), (E), (I), (M), and (N) show final decisions as a scatter plot in the *x*1*x*2-plane, where subjects’ actions are expected to cluster around the center of the workspace along each mini-block of 10 trials. (B), (F), (J), (N), and (R) show a histogram binning of the experimental scatter plots. (C), (G), (K), (O), and (S) present the change over the mean endpoint (averaged for both players) for each trial across the block of 40 trials. (D), (H), (L), (P), and (t) show the direction of adaptation in the endpoint space. The experimental data is shown in the top, the four continuous models are below.

The endpoint distribution of two coupled policy gradient learners performing in the continuous symmetric matching pennies game is shown in Figure 4E, and the corresponding version including intrinsic costs is shown in Figure 4I. In both cases the distribution of responses for both players is rather uniform. Figure 4M shows two coupled continuous Q-learning agents performing the same task, and Figure 4Q the same model including intrinsic costs. Contrarily to the gradient learning models, the Q-learning models manage to preserve substantial probability mass in the center of the workspace.

While all four models capture the effect of stable mean response frequencies—see Figures 4G,H, 4K,L, 4O,P, and 4S,T for comparison, the gradient learning models predict a more uniform distribution of responses, including in particular a high amount of extreme responses at the borders of the workspace that were mostly absent in the experiment. In contrast, the Q-learning models cluster in the center of the work space. Moreover, the inclusion of intrinsic costs that punish large deviations from behavior in the previous trial lead to a slightly stronger concentration of responses in the center of the work space in agreement with subject data. Altogether with the two-dimensional histogram binning of the scatter plots in Figures 4B, 4F, 4J, 4N, 4R the Q-learning model with intrinsic costs again achieves again the best fit to subjects’ behavior, confirmed by the lowest Euclidean distance between the fourth binned histogram—compare Supplementary Table 1.

As there is no change in response frequencies, assessing learning in this game is more challenging, even though the difference in endpoint distribution between the first 10 trials in a block and the last 10 trials was significant (KS-test, *p* > 0.01). To study the trial-by-trial adaptation process in more detail, we correlated subjects’ positional response from the current trial either to their positional response in the previous trial or the positional response of the other player in the previous trial. The correlations across four batches of ten consecutive trials that make up the blocks of 40 trials are shown in Figure 5. The positive correlations displayed by the solid lines indicate that subjects had a strong tendency to give the same kind of response in consecutive trials. In contrast, the close-to-zero correlations displayed by the dashed lines indicate that subjects’ behavior cannot be predicted from the other player’s response in the previous trial. These correlations were stable across the block of 40 trials. The close-to-zero correlations represented by the dashed lines are reproduced by all learning models. However, the solid lines representing the first-order autocorrelation behave quite differently in the models. The gradient learners start out with zero correlation between successive actions and increase this correlation slightly over trials, but always far below the high autocorrelation displayed by subjects. In contrast, the Q-learning models start out with an autocorrelation comparable in magnitude to the subject data, although the modeled autocorrelation decreases over time. In fact, in the limit of full convergence, the prediction is that the autocorrelation should vanish, thus, allowing convergence to full random play corresponding to the Nash equilibrium. While the subjects’ data does not show the predicted decrease in autocorrelation, the continuous Q-learning models fit the data best.

**Figure 5.**
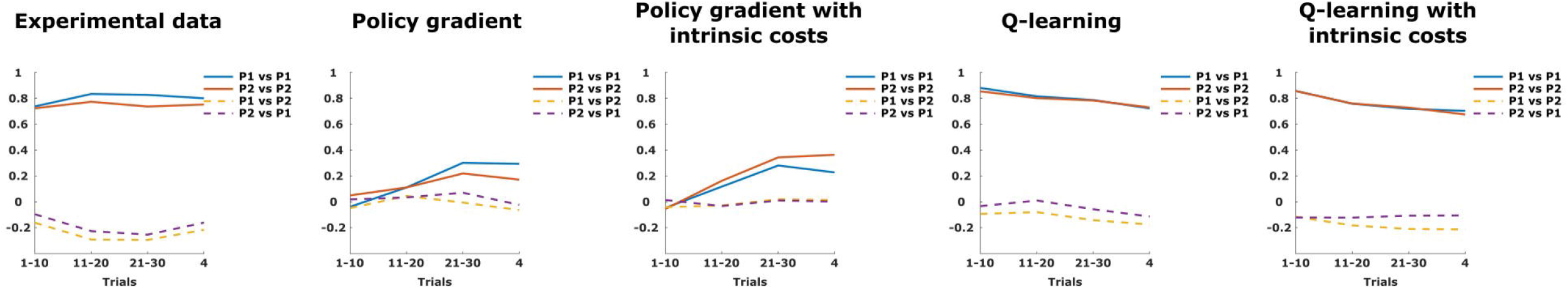
Autocorrelation in the symmetric matching pennies. The solid lines show the autocorrelation between actions from the current and the previous trial for both players. The dashed lines indicate cross-correlations between one players action in the current trial with another player’s action in the previous trial. From left to right: experimental data, policy gradient, policy gradient with intrinsic costs, Q-learning, Q-learning with intrinsic costs.

**Figure 6.**
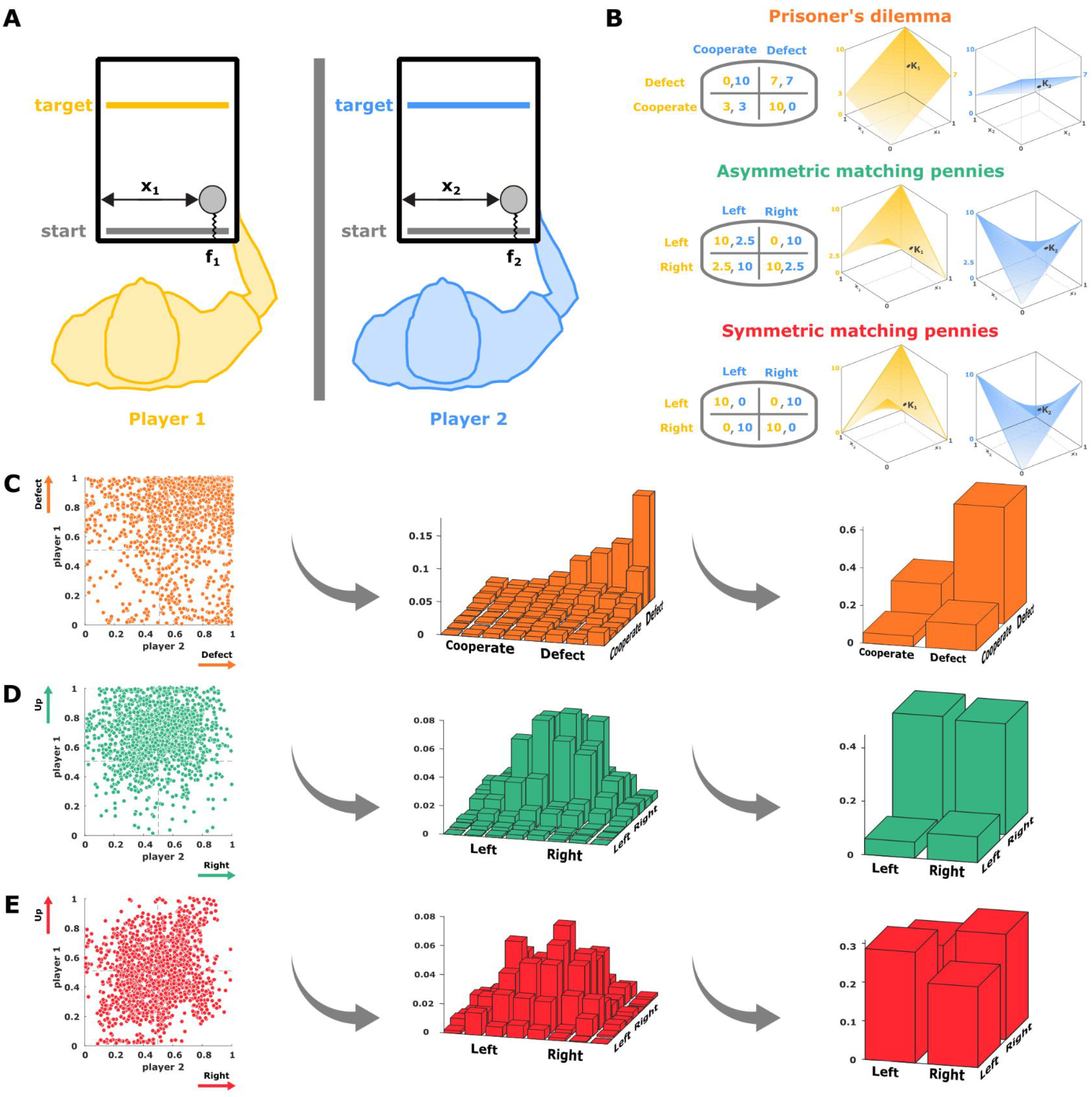
Experimental setup. **(A) Sensorimotor game.** Subjects were sitting next to each other without communication, and interacted through two handles that generated a force on their hand opposing their forward movement. **(B) Payoff matrices for each one of the 3 games.** 2-by-2 payoff matrix of player *i* determining the boundary values of the force payoffs at the extremes of the *x*1*x*2-space (left). Interpolation of the payoffs defining a continuous payoff landscape for a continuum of actions (right). **(C) Subjects results in the continuous Prisoners’ Dilemma game.** Final decisions shown as a scatter plot in the *x*1*x*2-space (left), where the single pure Nash equilibrium is located in the top-right corner at (1,1). The same responses binned into an 8-by-8 histogram (middle), and the corresponding categorical representation (right). **(D) Subjects results in the continuous Asymmetric Matching Pennies game.** Final decisions (left), where the mixed Nash equilibria is located at (0.8,0.5). The same responses binned into an 8-by-8 histogram (middle), and the corresponding categorical representation (right).**(E) Subjects results in the continuous Matching Pennies.** Final decisions game (left), where the mixed Nash equilibria is located at (0.5,0.5). The same responses binned into an 8-by-8 histogram (middle), and the corresponding categorical representation (right).

## Discussion

In this study we investigate human multi-agent learning in three different kinds of continuous sensorimotor games based on the three 2 × 2-matrix games of matching pennies, asymmetric matching pennies and the prisoners’ dilemma. In these games, subjects were haptically coupled and learned to move in a way that would reduce the experienced haptic coupling force resisting their forward motion, which served as a negative reward during the interaction. To explain subjects’ learning behavior across multiple repetitions, we compare their movements to different reinforcement learning models with or without internal costs arising when deviating from the previous action or average behavior. In previous studies^28^, the attainment of Nash equilibria in sensorimotor interactions has been reported, but without investigating the mechanisms of learning. In the current study we found, when simulating different learning models, that it is not enough to study binary discretizations of the continuous games as previously, as this coarse analysis does not reveal fine differences between learning models. However, clear differences between learning models arise when studying the games in their native continuous sensorimotor space. In particular, we found that a continuous Q-learning model with intrinsic action costs best explains subjects’ co-adaptation.

While we considered binary and continuous models in the paper to highlight the advantage the latter may have compared to the former, there is a gradual spectrum from discrete to continuous models. Naively, discrete models with more than two actions can be used to approximate the continuous case, but then learning is slower, because there are many more independent actions to learn. In the reinforcement learning literature^44^, this problem is remedied with multiple tilings, where each tiling contains a fixed number of tiles that cover the workspace. Having multiple tilings that are shifted with respect to each other introduces neighbourhood relationships where the number of tiles per tiling reflects the width of the neighbourhood. When formulating the Q-learning model in our examples as a discrete tiled agent, we get qualitatively similar results to the continuous agents, however, sometimes there are border effects. Therefore, we decided to use continuous models that more closely reflect the experimental setup, and that can be discretized for visualization.

In all of the games, the continuous Q-learning model trained without additional intrinsic costs tends to spread its probability fairly evenly across the workspace, and shows rather slow learning. The reason is not that Q-learning per se would be too slow to learn, but is rather a consequence of the fitting procedure that determines that the slow learning in that case provides the best fit, as faster learning would produce behavior that deviates from subjects’ behavior even more. However, we found that introducing an intrinsic cost that punishes large deviations from the previous action could alleviate this problem, allowing for more concentrated learning and making the Q-learning model qualitatively the best fit to human behavior. The intrinsic cost can be interpreted as a simple intrinsic cost for motor effort, that is not considered in the payoff function of the game, or it could be the consequence of an intrinsic motor planning cost that favors similar actions. As all our models were initialized with a uniform distribution, such an intrinsic cost could also be thought of as a non-uniform prior that one could use to initialize the models. Another alternative without explicitly initializing the priors this way, would be to conceptualize the minimization of the trial-by-trial deviation as minimizing the information difference between each action and the marginal distribution over actions given by the average behavior. Such an implicit information penalization of action policies can be regarded as a form of bounded rational decision-making, where decision-makers trade off between utility and information costs that are required to achieve a certain level of precision^45–47^. Such information costs have been previously suggested to model costs of motor planning and abstraction^48,49^. Another form of limited information-processing capability is subjects’ limited sensory precision regarding the perception of force, that we have modeled by assuming Gaussian sensory noise corresponding ten percent of the workspace^50,51^. Not including such sensory noise, predicts very fast convergence of the reinforcement learning models that is not observed in human subjects.

Compared to the Q-learning model, the policy gradient learners preferentially converged to strategies involving actions at the extremes, even though average behavior of these models fitted with game-theoretic predictions. This preference for extremes occurs not only for the particular parameterization that we chose (Kumaraswamy or beta-distribution), but also occurs for other parameterizations like the logit-normal model—compare Supplementary Figures 2-4. While this can only happen when the policy gradient models are initialized uniformly, a non-uniform initialization of these models must assign zero probability to the extremes of the workspace, and therefore trivially avoids this problem. When initialized uniformly, the gradient learners prefer the extreme solutions, as the reward gradient is steepest in this direction. This demonstrates that the choice of learning algorithm imposes a preference on the set of possible equilibrium solutions, that a priori would all be equally acceptable, due to the inherent learning dynamics. We conclude that it is important to study different learning algorithms for understanding sensorimotor interactions, as such behavior cannot be inferred from a game-theoretic analysis alone, that simply focuses on the Nash equilibrium concept. While our study can of course not exclude the possibility that there could be many other learning algorithms that could explain the data equally well or better, it shows also that it is not trivial that the Q-learning with intrinsic costs manages to capture the adaptive behavior of human interactions.

Previously, reinforcement learning models have been used to explain a host of phenomena in human strategic interactions including the emergence of dominant strategies^52,53^, conditional cooperation^54,55^, learning in extensive form games^56–58^, and transfer of reward-predictive representations^59,60^. The interactions in these studies were mostly based on cognitive strategies that participants could develop, as they were made explicitly aware of the meaning of different actions. Instead, we have focused on haptic interactions with forces that require implicit learning. The role of reinforcement learning in the context of implicit sensorimotor learning has also been previously examined^61–64^, including physical robot-human interactions^65,66^, however, not in the context of game theory and haptic interactions. Thus, our study adds to a growing body of research harnessing the power of reinforcement learning models to understand human interactions. While our task was simple enough to allow for model-free reinforcement learning, a challenge for the future remains to study model-based reinforcement learning in complex environments with multiple agents.

## Methods

### 1 Experimental methods

#### Participants

Sixteen naïve subjects participated in this study and provided written informed consent for participation. The study was approved by the ethics committee of Ulm University. All sixteen participants were undergraduate students that were compensated for their time with an hourly payment of 10 euros, where they engaged in sensorimotor interactions equivalent to two versions of the matching pennies game. The data of another sixteen subjects was reanalyzed from a previous study^28^, where they engaged in sensorimotor interactions equivalent to the prisoners’ dilemma under the same kind of experimental conditions presented here in the current study. All methods were carried out in accordance with relevant guidelines and regulations.

#### Setup

The experiment was run on two vBOTs haptic devices^67^ that were connected to a virtual reality environment in which two subjects could control one cursor each by moving a handle in the horizontal plane. This setup provides both subjects separately with a visual feedback of their own cursor position, without showing a direct view of their hand or the hand of the other player—see Figure 6A. The subjects were haptically coupled by applying individual forces on the two handles representing a negative payoff. Both the forces and the cursors’ positions were updated and recorded with a sampling frequency of 1 kHz throughout each trial.

#### Experimental design

To begin each trial, subjects had to simultaneously locate their cursor on the 15-cm-wide starting bar placed at each bottom side of their respective workspace. Each participant’s workspace was constrained by the vBOT simulating solid walls at the extremes left (−8*cm*) and right (+8*cm*) for both players. A beep sound informed players that a valid starting position was chosen and a 15-cm-wide target bar was displayed in front of each player with the same distance that was randomly drawn from the uniform distribution between 5 and 20 cm—see Figure 6A. Subjects were instructed to move as soon as they heard the beep. Each participant’s goal was to hit their respective target bar with their cursor, for which they had to move forward (y-direction) and touch the bar within a maximum time period of 1500 ms. Each trial was finished as soon as both players had hit their respective target bar, which they could touch anywhere along its length. For the analysis, we interpreted this horizontal position of the target crossing as subjects’ final choice.

To determine the payoffs of the interaction, each player’s cursor was attached to a simulated one-dimensional spring whose equilibrium point was located on the starting bar. This generated a force *F_i_* on their hand in the negative y-direction that opposed their forward movement with *F_i_* = –*K_i_y_i_*, where *y_i_* corresponds to the y-distance of the cursor of player *i* ∈ {1,2} from the starting bar. Crucially, the spring constants *K_i_* were not constant but variable functions *K_i_* (*x*_1_, *x*_2_) depending on the normalized horizontal positions *x*_1_ and *x*_2_ of both participants, where *x_i_* = 0 corresponds to extreme left and *x_i_* = 1 represents extreme right of the workspace of player *i*. In particular, the spring functions *K_i_*(*x*_1_, *x*_2_) were determined by the bilinear interpolation

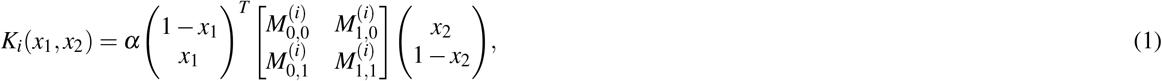

where *α* was a constant scaling parameter set to 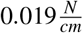 throughout the experiment, and the 2-by-2 matrices *M*^(*i*)^ corresponds to the 2-by-2 payoff matrix of player *i* and determine the boundary values of the force payoffs at the extremes of the *x*_1_*x*_2_-space. Consequently, 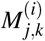 is the payoff assigned to player *i* when player 1 takes action *x*_1_ = *k* and player 2 takes action *x*_2_ = 1 – *j* where *j,k* ∈ {0,1}. Each one of the three games analyzed in this study was defined by a different set of payoff matrices corresponding to the 2-by-2 games of the prisoner’s dilemma, asymmetric matching pennies, and symmetric matching pennies—see Figure 6B. Players’ actions are considered as the final positions in the *x*1*x*2-space, where the continuous responses can be categorized by using different binning (see Figures 6C,D,E).

To encourage learning, the payoff matrices were permuted every 40 trials, so that the mapping between action and payoff had to be relearned every 40 trials. There were four possible permutations corresponding to the possible reassignments *x*_1_ → (1 – *x*_1_) and *x*_2_ → (1 – *x*_2_). Subject pairs repeated 20 blocks of 40 trials each, totaling in 800 trials over the whole experiment. For the purpose of analysis, we took subjects’ Before the experiment, participants were instructed to bring the cursor to the target bar while trying to minimize the resistive forces experienced during the movement. In addition, subjects were not allowed to communicate with each other during the experiment.

#### Prisoner’s dilemma motor game

In the pen-and-paper version of the prisoners’ dilemma the 2-by-2 payoff matrices are given by

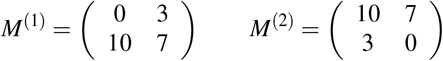

and represent a scenario where two delinquents are caught separately by the police and face the choice of either admitting their crime (cooperating) or denying it (defecting), while being individually interrogated with no communication allowed. For each player it is always best to defect, no matter what the other player is doing, even though cooperation would lead to a more favorable outcome if both players choose to do so. Hence, there is only one pure Nash equilibrium given by defect/defect. In our continuous sensorimotor version of the game, the Nash equilibrium corresponds to a corner of the *x*_1_*x*_2_-space—see Figure 6C.

#### Matching pennies motor games

In the classic version of the symmetric matching pennies game the 2-by-2 payoff matrices are given by

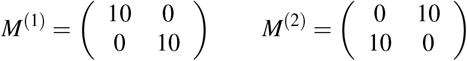

and can be motivated from the scenario of 2 football players during a penalty kick with the two actions *left* and *right*. In this case player 2 (e.g. the goalkeeper) will try to select the matching action (e.g. *x*_1_ = *right*/*x*_2_ = *right*) to stop the ball, while player 1 (e.g. the striker) will try to unmatch by aiming the shot away from player 1 (e.g. *x*_1_ = *left*/*x*_2_ = *right*). Choosing an action deterministically in this game is a bad strategy, since the opponent can exploit predictable behavior. The mixed Nash equilibrium is therefore given by a pair of distributions (*p*_1_;*p*_2_) with *p*_1_(*left*) = *p*_1_(*right*) = 0.5 and *p*_2_ (*left*) = *p*_2_(*right*) = 0.5 (see Figure 6E). In the continuous sensorimotor version of the experiment there are infinitely many mixed Nash equilibria corresponding to the set of all distributions {*p*_1_,*p*_2_} where 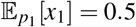 and 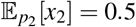 assuming *x_i_* ∈ [0; 1]—see Supplementary Material.

In the 2-by-2 version of the asymmetric matching pennies game the payoff matrices are given by

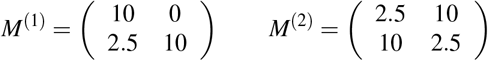

which is similar to the symmetric scenario, only that one player now has a higher incentive to choose a particular action (e.g. the striker gets awarded more points when choosing *x*_2_ = *left*. The mixed Nash equilibrium is then given by a pair of distributions (*p*_1_;*p*_2_) with *p*_1_(*left*) = 0.8, *p*_1_(*right*) = 0.2 and *p*_2_(*left*) = *p*_2_(*right*) = 0.5 (see Figure 6D.). In the continuous sensorimotor version of the experiment there are infinitely many mixed Nash equilibria corresponding to the set of all distributions {*p*_1_, *p*_2_} where 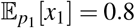 and 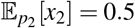 assuming *x_i_* ∈ [0; 1]—see Supplementary Material.

### 2 Theoretical methods

#### Computational models

##### Structure

Every model consists of two artificial agents whose parameters are updated over the course of 40 trials. Like in the actual experiment, every agent decides independently from the other agent about an action, observes the reward signal in the form of a punishing force and adapts his strategy. The models can be categorized in binary action models with action space 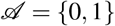, and continuous action models that operate on the action space 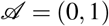. All models follow the same pattern:

- At the beginning of every trial *t* every agent *i* samples an action 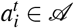 from the distribution 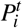.
- Both agents then take their respective action simultaneously and receive a reward/punishment signal in the form of a force 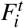. This force is calculated as a continuous linear interpolation of the reward that would be assigned to actions 0 and 1 according to the payoff matrix *M*.
- Depending on the action and reward, the agents adapt their strategy 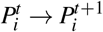 to optimise future rewards.

##### Binary action models

Binary action models are limited to the actions 0 and 1 meaning that they play the cognitive rather than the sensorimotor version of the different games.

###### Model 1: Q-Learning

As in standard Q-Learning, agents using the first model represent their knowledge in the form of Q-value lookup table. Since there are only two actions the lookup table for player i has two values, namely:

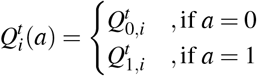

For the distribution 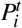 we chose a softmax-distribution with parameter ***β**_i_*:

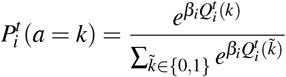

and the updates for the Q-values are of the following form:

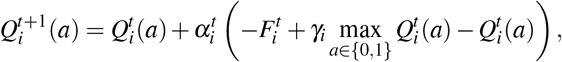

where 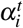 is a learning rate that decreases exponentially over the course of the trials. All Q-values are initialized as 0, such that the agents start the game without any initial knowledge.

###### Model 2: Gradient Decent

The gradient decent model learns a parameterized policy that can select actions without consulting a value function or requires a maximization over many samples. The policy used by the binary gradient decent learner is the Bernoulli probabilty distribution with outcomes {0,1}. An action *a* is determined by sampling from the policy:

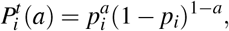

where 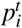 is a parameter that is adapted whenever a player receives reward or punishment in form of a force 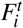 for their current action 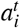.

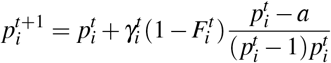

with an exponentially decaying learning rate parameter *γ* that is fitted to the behaviour of the human participants. The parameter is initialized as 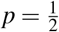 such that both actions are equally probable for the initial policy.

##### Continuous action models

Continuous models are closest to the sensorimotor version of the game in that they are able to sample actions from the full interval (0, 1).

###### Model 3: Gradient Descent

The gradient decent model directly learns a parameterized policy that can select actions without consulting a value function. The parameterized policy used in this study is given by the Kumaraswamy probability distribution on the interval [0,1]. An action a is determined by sampling from the policy:

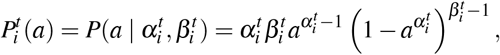

where 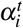 and 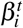 are shape-parameters that are adapted whenever a player receives reward or punishment in form of a force for their current action. For an action 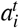 and punishing force 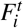, the parameters are updated according to the following update equations:

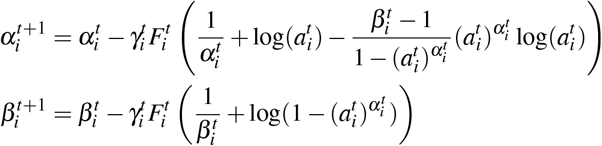

with an exponentially decaying learning rate parameter γ that is fitted to the behaviour of the human participants. The shape-parameters are initialized as *α* = *β* = 1 such that the initial policy for all players is the uniform distribution on (0,1).

###### Model 4: Q-Learning with Gaussian basis functions

This model is an extension of the discrete Q-Learning model to the continuous action space (0,1). Its core consists of an action value function with radial basis functions such that the *real* action value function 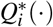 is well reflected by a set of weights 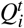 that are adapted throughout the trials *t* = 1,2,… and a set of parameters (*c_i_, **σ**_i_*) that are held constant. For the simulation we chose Gaussian basis functions centered around *c_k_* with standard deviation ***σ**_k_* such that we can generate action samples by drawing them from a Gaussian mixture model.

So, in order to generate an action 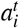 at time *t*, we sample an index *j* from an categorical distribution with parameters 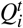 and then draw a sample *a* from the *j*-th basis function in the Gaussian mixture. Samples that lie outside of the interval (0,1) are rejected and we sample again.

Unlike the discrete case we update all weights *Q* in every trial according to the following rule. If *a_t_* and *F_t_* are the action taken and the punishing force at time *t*, then

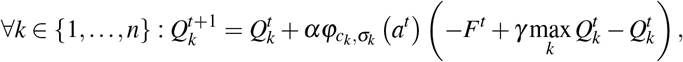

where *α* is a learning rate parameter and *γ* is a discount factor. All weights were initialized as 0.

##### Optimizing parameters

All parameters were optimized such that the simulated agents’ behavior fitted the behavior of the average player in a specific player slot (i.e. player 1 or player 2). For the optimization we minimized the absolute error:

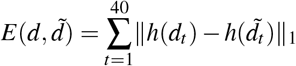

that occurred over 40 trials between the participants data *d* and the simulation’s results 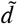. The histogram function *h* maps the data points onto an 8 × 8 matrix where each entry of this matrix contains the normalized amount of data points that lie in the corresponding bin of an 8 × 8 binned histogram. The time index *t* of *d_t_* indicates that all data points from the first trial up until trial *t* are used to generate the matrix.

## Acknowledgments

This study was funded by European Research Council (ERC-StG-2015—ERC Starting Grant, Project ID: 678082, BRISC: Bounded Rationality in Sensorimotor Coordination).

## Author contributions

CLL, GS, and DB designed the experiment. CLL performed experiments and analyzed the data. CLL and GS generated predictions from computer simulations. DB supervised the project. CLL and DB wrote the paper.

