## Supplementary figures for "Nash equilibria in human sensorimotor interactions explained by Q-Learning"

### Nash equilibria in the continuous matching pennies games

Let the actions of player 1 and player 2 be two independent random variables $X_1$ and $X_2$ on $\Omega = [0, 1]$ and let $M \in \mathbb{R}^{2\times 2}$ be the payoff matrix for the continuous (asymmetric) matching pennies game, i.e.

$$(M_{i,j,1}) = \begin{pmatrix} a & 0 \\ 0 & 1 \end{pmatrix}$$
$$(M_{i,j,2}) = \begin{pmatrix} 0 & 1 \\ 1 & 0 \end{pmatrix}$$

where $a$ is a constant. Then every pair of proper probability distributions of $X_1$ and $X_2$ such that $\mathbb{E}[X_1] = \frac{1}{2}$ and $\mathbb{E}[X_2] = \frac{1}{a+1}$ is a Nash equilibrium.

*Proof.* The continuous payoff $U_k$ for player $k$ is given by the payoff interpolation

$$U_k(x_1, x_2) = (1 - x_1)\left(x_2 M_{1,1,k} + (1 - x_2)M_{1,2,k}\right) + x_1\left(x_2 M_{2,1,k} + (1 - x_2)M_{2,2,k}\right),$$

Then the expected payoff of player $k$ is

$$\mathbb{E}[U_k] = \int_{\Omega^2} p(X_1 = x_1, X_2 = x_2)U_k(x_1, x_2)dx,$$

where $p$ is the joint distribution of $X_1$ and $X_2$. Since $X_1$ and $X_2$ are assumed to be independent $p$ factorizes into $p(X_1 = x_1, X_2 = x_2) = p_1(x_1)p_2(x_2)$ and it follows that

$$\begin{aligned}
\mathbb{E}[U_1] &= \int_\Omega \int_\Omega p_1(x_1)p_2(x_2)\left((1-x_1)(x_2 a) + x_1(1-x_2)\right)\, dx_2\, dx_1 \\
&= \int_\Omega p_1(x_1)\left(\int_\Omega p_2(x_2)x_1\, dx_2 + \int_\Omega p_2(x_2)x_2 a\, dx_2 - \int_\Omega p_2(x_2)(1+a)x_1x_2\, dx_2\right)\, dx_1 \\
&= \int_\Omega p_1(x_1)\left(x_1 + a\mathbb{E}[X_2] - (1+a)x_1\mathbb{E}[X_2]\right)\, dx_1 \\
&= a\mathbb{E}[X_2] + \int_\Omega p_1(x_1)(1-(1+a)\mathbb{E}[X_2])x_1\, dx_1,
\end{aligned}$$

which is independent of $p_1(x_1)$ whenever $\mathbb{E}[X_2] = \frac{1}{1+a}$. In that case player 1 is indifferent about the distribution of his actions, making it a Nash-strategy for player 2.

Similarly, it is

$$\begin{aligned}
\mathbb{E}[U_2] &= \int_\Omega \int_\Omega p_1(x_1)p_2(x_2)\left((1-x_1)(1-x_2) + x_1x_2\right)\, dx_2\, dx_1 \\
&= \int_\Omega p_2(x_2)\left(\int_\Omega p_1(x_1)\, dx_1 - \int_\Omega p_1(x_1)x_1\, dx_1 - \int_\Omega p_1(x_1)x_2\, dx + 2\int_\Omega p(x_1)x_1x_2\, dx_1\right)\, dx_2 \\
&= (1-\mathbb{E}[X_1]) + \int_\Omega p_2(x_2)(2\mathbb{E}[X_1]-1)x_2\, dx_2,
\end{aligned}$$

which is independent of $p_2(x_2)$ whenever $\mathbb{E}[X_1] = \frac{1}{2}$. □

### Categorical Analysis

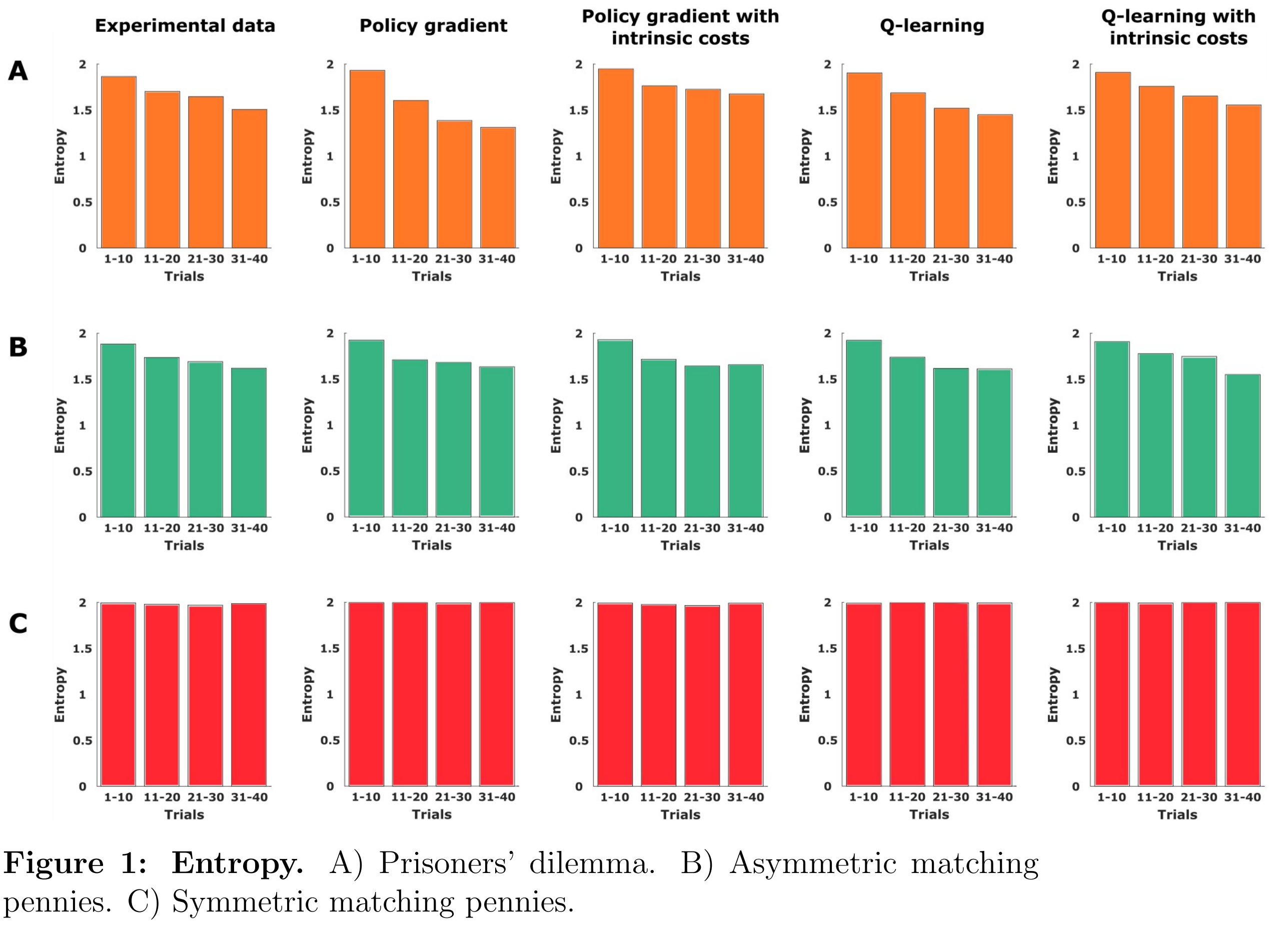


**Figure 1: Entropy.** A) Prisoners' dilemma. B) Asymmetric matching pennies. C) Symmetric matching pennies.

### Continuous Analysis

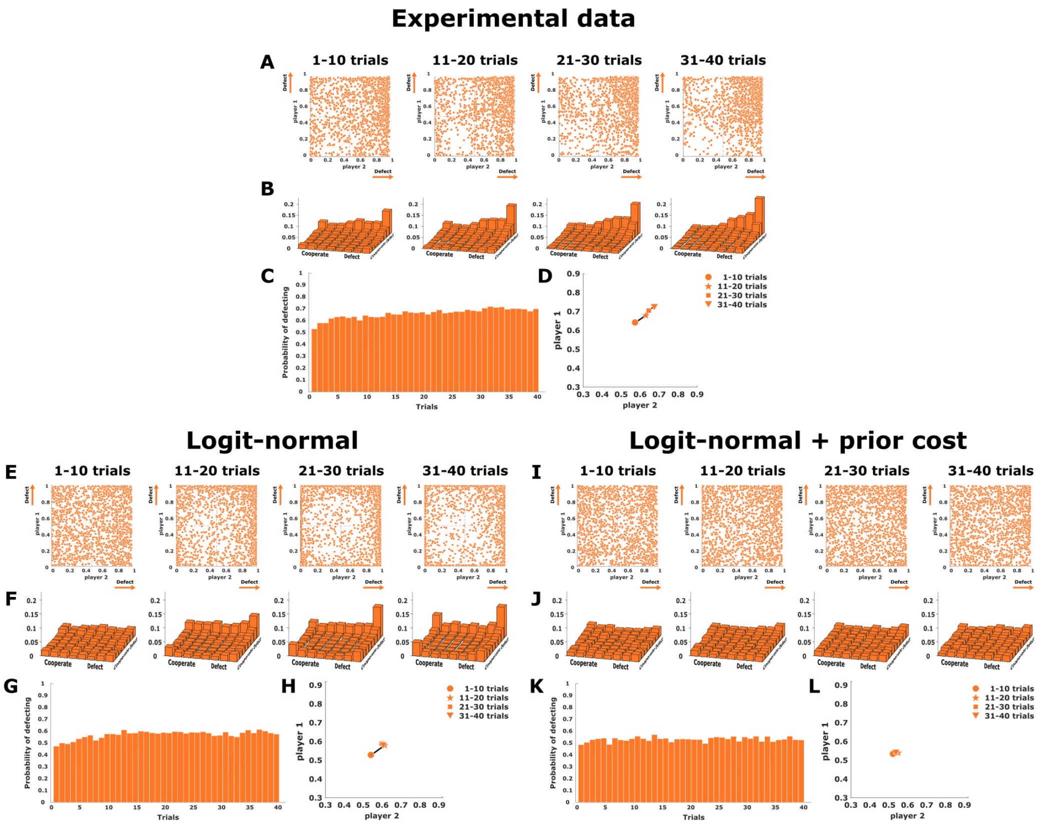


**Figure 2: Prisoner's dilemma, logit normal.** (A), (E), and (I) show scatter plots of final decisions in the $x1x2$-plane, where subjects' actions are expected to cluster around the single pure Nash equilibrium located in the top-right corner at position (1,1). (B), (F), and (J) show two-dimensional histograms binning the experimental scatter plots. (C), (G), and (K) represent the change of the mean endpoints (averaged for both players) for each trial across the block of 40 trials. (D), (H), and (L) show the direction of adaptation in the endpoint space. The experimental data is shown at the top, the two continuous models are shown below.

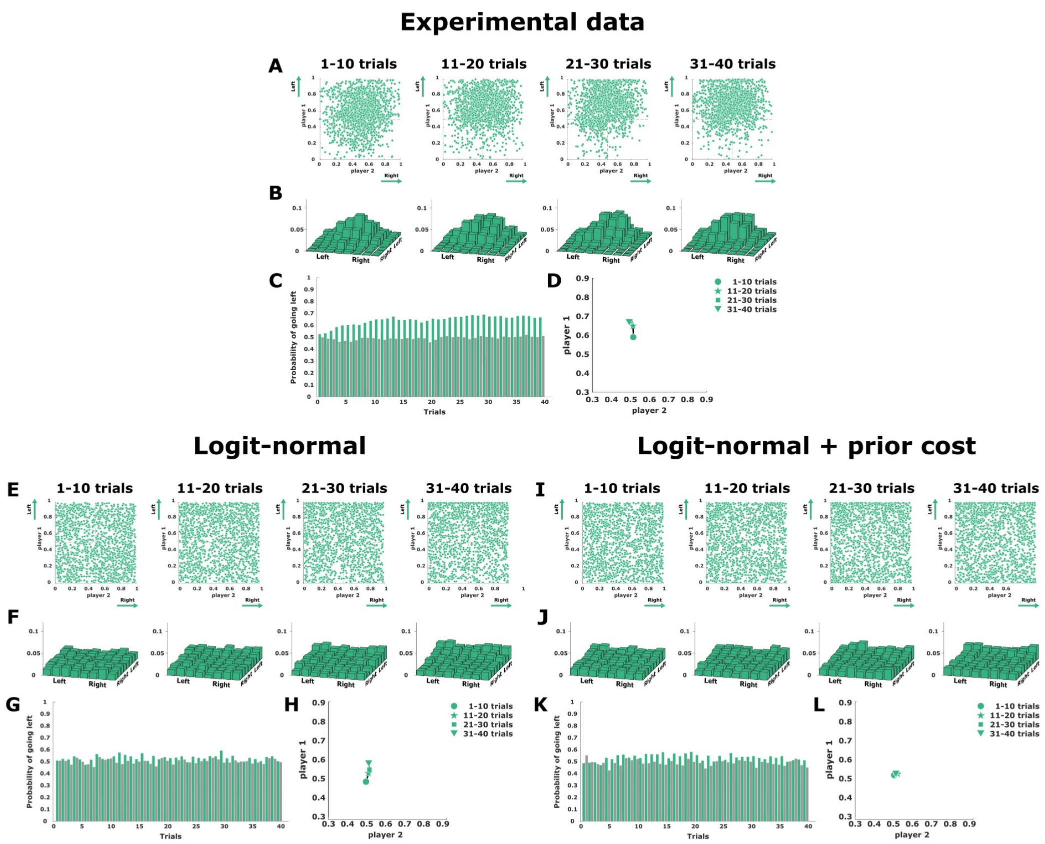


**Figure 3: Asymmetric matching pennies, logit normal.** (A), (E), and (I) show final decisions as a scatter plot in the $x1x2$-plane, where subjects' actions are expected to cluster in top quadrants along each mini-block of 10 trials. (B), (F), and (J) show a two-dimensional histogram binning of the experimental scatter plots. (C), (G), and (K) present the change over the mean endpoint (averaged for both players) for each trial across the block of 40 trials. (D), (H), and (L) show the direction of adaptation in the endpoint space. The experimental data is shown on the top, the two continuous models are below.

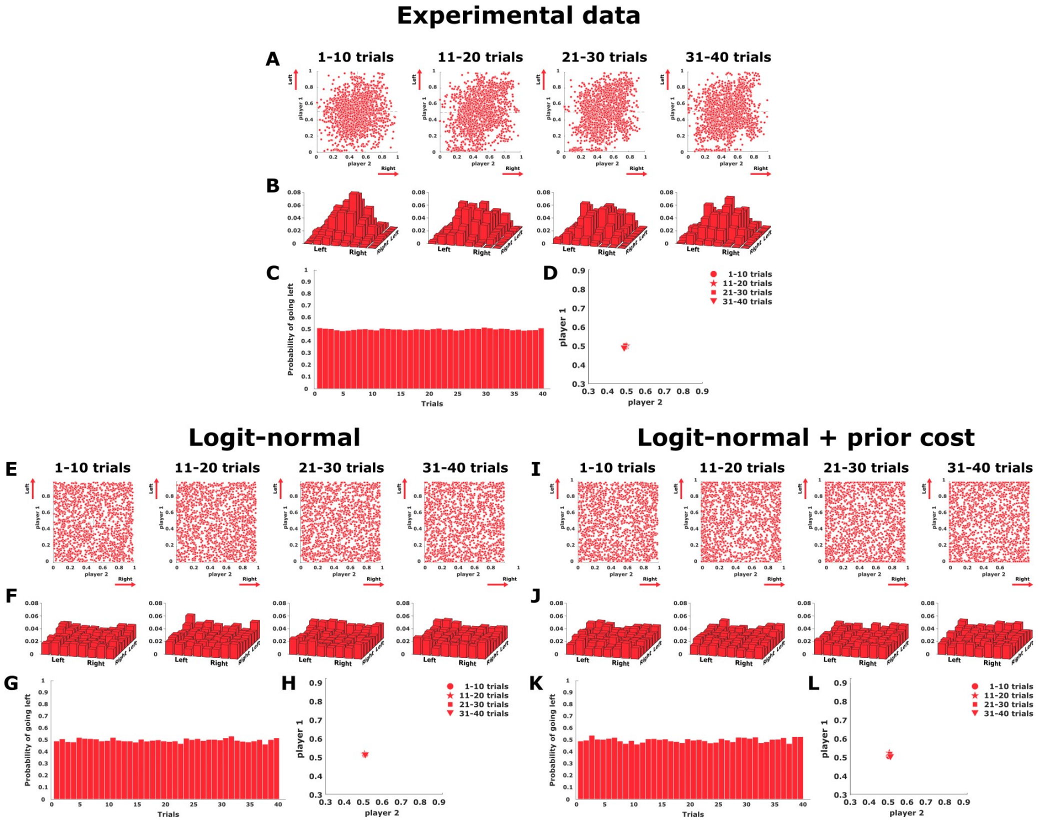


**Figure 4: Symmetric matching pennies, logit normal.** (A), (E), and (I) show final decisions as a scatter plot in the $x1x2$-plane, where subjects' actions are expected to cluster around the center of the workspace along each mini-block of 10 trials. (B), (F), and (J) show a histogram binning of the experimental scatter plots. (C), (G), and (K) present the change over the mean endpoint (averaged for both players) for each trial across the block of 40 trials. (D), (H), and (L) show the direction of adaptation in the endpoint space. The experimental data is shown in the top, the two continuous models are below.

| Game | Euclidean distance | | | | Mean-squared error | | | |
|---|---|---|---|---|---|---|---|---|
| | PG | PG+pc | Q | Q+pc | PG | PG+pc | Q | Q+p[illegible] |
| Prisoners' dilemma | 0.12 | 0.1 | 0.14 | 0.08 | 0.003 | 0.005 | 0.014 | 0.00[illegible] |
| Asymmetric MP | 0.17 | 0.19 | 0.12 | 0.1 | 0.003 | 0.005 | 0.001 | 0.00[illegible] |
| Symmetric MP | 0.15 | 0.16 | 0.08 | 0.08 | $0.34e-3$ | $0.34e-3$ | $0.19e-3$ | $0.17e$ [illegible] |

**Table 1:** Evaluation.
